# The integrated stress response regulates BMP signaling through effects on translation

**DOI:** 10.1101/266189

**Authors:** Elke Malzer, Caia S. Dominicus, Joseph E. Chambers, Souradip Mookerjee, Stefan J. Marciniak

**Affiliations:** Cambridge Institute for Medical Research (CIMR), University of Cambridge Wellcome Trust/MRC Building, Hills Road, Cambridge, CB2 0XY, UK.; Department of Medicine, University of Cambridge, Addenbrooke’s Hospital, Hills Rd, Cambridge CB2 0SP

## Abstract

Developmental pathways must be responsive to the environment. Phosphorylation of eIF2α enables a family of stress sensing kinases to trigger the integrated stress response (ISR), which has pro-survival and developmental consequences. Mutations of the ISR kinase GCN2 have been implicated in the development of pulmonary arterial hypertension, a disorder known to be associated with defects of BMP signaling, but how the ISR and BMP signaling might interact is unknown. Here we show in *Drosophila* that GCN2 antagonises BMP signaling through direct effects on translation and indirectly via the transcription factor *crc* (dATF4). Expression of a constitutively active GCN2 or loss of the eIF2α phosphatase dPPP1R15 impair developmental BMP signaling in flies. In cells, inhibition of translation by GCN2 blocks downstream BMP signaling. Moreover, loss of d4E-BP, a target of *crc*, augments BMP signaling *in vitro* and rescues tissue development *in vivo*. These results identify a novel mechanism by which the ISR modulates BMP signaling during development. Since abnormalities of both GCN2 and BMP signaling lead to pulmonary hypertension, these findings may have wider relevance for the development of therapies for this disease.

## Introduction

Pulmonary arterial hypertension (PAH) is a family of diseases that predominantly affects young adults and carries a high mortality. Although most cases are idiopathic, in 70% of familial cases and 20% of sporadic cases heterozygous germline mutations are identified in the type II BMP receptor (*BMPR2*) (Machado et al., 2001, International et al., 2000, Thomson et al., 2000). The penetrance of *BMPR2* mutation is highly variable suggesting that additional modifying factors must exist. Recently, two rare subtypes of PAH, pulmonary veno-occlusive disease (PVOD) and capillary haemangiomatosis, were shown to be caused by mutations of *EIF2AK4*, which encodes the kinase GCN2 (Eyries et al., 2014, Best et al., 2014). Interestingly, *BMPR2* mutations have also been associated with PVOD, suggesting that similar mechanisms may underlie typical PAH and PVOD (Runo et al., 2003, Montani et al., 2008).

GCN2 belongs to a family of stress-sensing kinases that phosphorylate the alpha subunit of eukaryotic translation initiation factor 2 (eIF2α) to activate the Integrated Stress Response (ISR) (Harding et al., 2003). When eIF2α is phosphorylated, the translation of most mRNAs is reduced to limit amino acid consumption; however, a small subset is translated more efficiently, including the mRNA encoding the transcription factor ATF4 (Vattem and Wek, 2004, Lu et al., 2004). Targets of ATF4 aid survival by promoting amino acid import and the biosynthesis of aminoacyl-tRNAs (Harding et al., 2003). One ISR target gene encodes an eIF2α phosphatase called PPP1R15A (also called GADD34), which dephosphorylates eIF2α to restore protein synthesis and permit the translation of ISR targets (Novoa et al., 2001, Marciniak et al., 2004, Ma and Hendershot, 2003).

The importance of the ISR during stress is well appreciated, but it also plays a less well-understood role during development. In mice, a lack of the ISR owing to mutation of eIF2α (eIF2α^S51A^) causes growth retardation *in utero* and perinatal death (Scheuner et al., 2001), while exaggeration of the ISR by deleting both eIF2α phosphatases (PPP1R15A & B) causes very early embryonic death (Harding et al., 2009). Mutation of the ISR kinase PERK in humans and mice has multiple effects on development including skeletal dysplasia (Delepine et al., 2000). At least some of the developmental effects of the ISR are mediated by ATF4. Consequently, *Atf4*^-/-^ mice have impaired osteoblast differentiation and bone mineralisation (Yang et al., 2004). We previously showed that ATF4 regulates protein secretion via the transcription factor CHOP (Marciniak et al., 2004) and that *Chop*^-/-^ mice have retarded bone formation (Pereira et al., 2006). The role of the ISR in osteogenesis may involve bidirectional crosstalk between eIF2α phosphorylation and BMP signaling. For example, treatment of primary bone cultures with BMP2 triggers endoplasmic reticulum stress and induces ATF4 in a PERK-dependent manner (Saito et al., 2011), while CHOP promotes differentiation of osteoblasts upon treatment with BMP (Shirakawa et al., 2006).

How BMP and GCN2 signaling might interact is not known. Here, we use *Drosophila melanogaster* to identify a novel mechanism by which GCN2 regulates BMP-dependent MAD phosphorylation.

## Results

### Depletion of dPPP1R15 or dGCN2 alters wing venation

To understand the role of the ISR in tissue development, we used the model organism *Drosophila melanogaster*. It shares ISR components with mammals (Malzer et al., 2010, Malzer et al., 2013), but its smaller genome reduces redundancy. We previously reported that changes in the expression of the eIF2α kinase dGCN2 or the eIF2α phosphatase dPPP1R15 impair fly development (Malzer et al., 2013). To determine which tissues are sensitive to altered ISR signaling, we now expressed *ppp1r15* RNAi under the control of a panel of tissue-selective drivers (Supplementary Fig 1A). Ubiquitous knockdown of *ppp1r15* or knockdown limited to the ectoderm markedly impaired larval development. In contrast, *ppp1r15* depletion in multiple tissues including the fat body, somatic muscle, salivary gland, midgut visceral mesoderm, eye, central nervous system (CNS), ring gland, or heart had no detectable consequence for development. However, the use of the *escargot* driver *(esgGAL4)*, which is expressed in several tissues including the imaginal discs, caused larval delay at the third instar stage (Supplementary Fig 1B-D). Larvae expressing *esgGAL4* driven *ppp1r15* RNAi (*esg>ppp1r15* RNAi) were followed until 21 days after egg laying (AEL) with less than 10% reaching adulthood. Similarly, using an engrailed driver (*enGAL4)* to express *ppp1r15* RNAi primarily in the posterior compartments of the imaginal discs also led to developmental delay (Supplementary Fig 1E). Larvae expressing *enGAL4* driven *ppp1r15* RNAi (en>*ppp1r15* RNAi) were delayed, but approximately 45% reached adulthood by 14 days. Delayed larvae appeared phenotypically normal, continuing to feed and grow in size.

Since loss of the phosphatase dPPP1R15 would be expected to cause hyper-phosphorylation of its substrate eIF2α, we hypothesized that loss the eIF2α kinases might rescue the effects of *ppp1r15* RNAi. Indeed, depletion of the eIF2α kinase *perk* by RNAi driven either by *esgGAL4* or *enGAL4* largely rescued *ppp1r15* RNAi expressing animals to adulthood (Supplementary Fig 1D-1E). Similarly, although depletion of *gcn2* by RNAi driven either by *esgGAL4* or *enGAL4* caused a modest developmental delay, with only ≈70% of animals reaching adulthood by 14 days, *esgGAL4>gcn2* RNAi partially rescued the developmental delay caused by *ppp1r15* knockdown (Supplementary Fig 1C-E).

These results revealed that the development of *Drosophila* could be impaired by a genetic perturbation predicted to enhance phosphorylation of eIF2α. This sensitivity displayed a restricted tissue distribution that included the imaginal discs, but excluded much of the animals’ tissue mass. Raising animals on high protein diet rather than standard food had no measureable effect on the frequencies of wing phenotypes nor on the number of animals eclosing (not shown). Low protein diets led to fewer adults, but the frequencies of each phenotype was unaffected. These findings suggested that protein deprivation was unlikely to account for the observed role of the ISR in our model.

In most respects, *en>ppp1r15* RNAi animals appeared normal, although their wings lacked the anterior cross vein (ACV) (Fig 1A, open triangle). By contrast, depletion of dGCN2 in the posterior compartment of the wing (*en>gcn2* RNAi) led to ectopic venation between longitudinal veins 4 (L4) and L5 (Fig 1A, closed triangles). Frequently, *en>gcn2* RNAi animals lacked the posterior half of the ACV (Fig 1A&B). When *en>ppp1r15* RNAi and *en>gcn2* RNAi were expressed together, the phenotype more closely resembled that of *en>gcn2* RNAi with ectopic venation between L4 and L5, and frequent absence of the posterior portion of the ACV (Fig 1A&B). The effect of depleting dPPP1R15 on venation appeared to be dose-dependent, since augmentation of RNA interference by co-expression of dicer2 led to combined loss of the ACV, the posterior cross vein (PCV) and L4 (Fig 1C).

**Figure 1.**
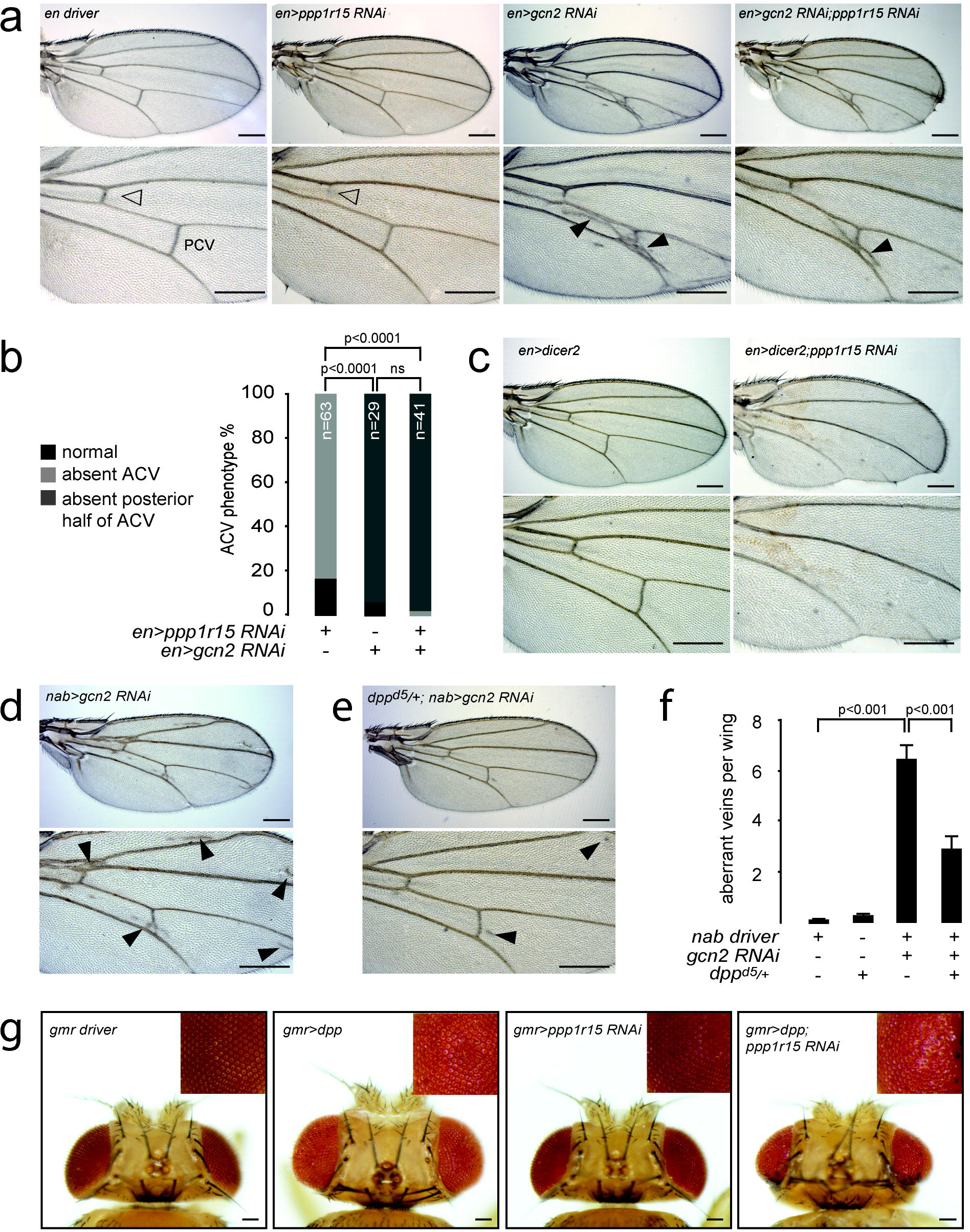
Depletion of dPPP1R15 or dGCN2 alters wing venation. (A) Representative photomicrographs (5x objective) of adult wings of the indicated genotypes. Lower panels are enlargements of the crossvein territories: anterior crossvein (open arrowhead) and posterior crossvein (PCV). Note extra venation (closed arrowheads) in wings expressing *gcn2* RNAi. Scale bars = 250µm. (B) Quantification of ACV phenotypes. For brevity, *enGAL4>UAS-ppp1r15* RNAi is indicated as *en>ppp1r15* RNAi. *enGAL4>UAS-gcn2* RNAi is indicated as *en>gcn2* RNAi. “n” denotes number of animals counted. P values calculated using ^2^ statistics with Bonferroni correction for multiple comparisons. (C) Representative photomicrographs of adult wings (5x objective) of the indicated genotypes. *en>dicer2* indicates *enGAL4>UAS-dicer2*. *en>dicer2;ppp1r15* RNAi indicates *enGAL4>UAS-dicer2;ppp1r15* RNAi. Lower panels are enlargements of the crossvein territories. Scale bars = 250µm. (D-E) Representative photomicrographs of adult wings (5x objective) of the indicated genotypes. *nab>gcn2* RNAi indicates *enGAL4>UAS-gcn2* RNAi. *dpp^d5/+^; nab>gcn2* RNAi indicates *Dpp^d5/+^; nab>UAS-gcn2* RNAi. Lower panels are enlargements of the crossvein territories. Note extra venation (closed arrowheads). (F) Quantification of wings from (D) and (E). Scale bars = 250µm. (G) Representative photomicrographs of adult eyes (dorsal view) of the indicated genotypes, insert shows zoom of eye. Scale bar = 200µm.

When *gcn2* RNAi was driven by *nabGAL4*, ectopic venation was observed adjacent to the longitudinal veins (Fig 1D&F, closed triangles). Because cross vein formation is sensitive to dpp (*Drosophila* BMP2/4) signaling (Ray and Wharton, 2001), we examined the effect of manipulating dGCN2 and dPPP1R15 in animals with one hypomorphic allele of *dpp*, *dpp^d5^* (Capdevila et al., 1994). *dpp^d5/+^* heterozygous animals retained normal wing venation (Supplementary Fig 1F) while *dpp^d5/+^* animals showed significantly less ectopic venation caused by depleting dGCN2 with *nab>gcn2* RNAi (Fig 1DF). In contrast, loss of one wild type allele (*dpp^d5/+^*) sensitized animals to depletion of *ppp1r15*, causing loss of posterior wing blade tissue and distal portions of L5 (Supplementary Fig 1F).

These results suggested that components of the ISR, specifically dGCN2 and dPPP1R15, could modulate wing imaginal disc development and that this might involve effects on dpp/BMP signaling. In support of this, we also observed that depletion of dally, a cell-surface glypican involved in dpp signaling (Belenkaya et al., 2004), also interacted genetically with dPPP1R15 and dGCN2. Alone, expression of *dally* RNAi using the nab driver had no effect on wing venation, but when combined with knockdown of *ppp1r15* it exacerbated the loss of wing blade tissue and, once again, led to loss of distal portions of L5 (Supplementary Fig 1F). When combined with *nab*>*gcn2* RNAi, depletion of *dally* caused disorganised venation (not shown).

Overgrowth of the eye reports on elevated dpp signaling (Li and Li, 2006). We therefore tested the effect of depleting dPPP1R15 in the eye using a *gmrGAL4* driver (Fig 1G). As expected, overexpression of dpp in the eye led to eye overgrowth. Knockdown of dPPP1R15 alone had no detectable effect on eye development, but when combined with overexpression of dpp, it rescued eye growth to a normal size, albeit with a rough eye phenotype.

These observations suggested that the developmental effects of modulating the ISR were sensitive to the intensity of dpp signaling, revealing a novel genetic interaction between the ISR and BMP pathways during fly development.

### dPPP1R15 or dGCN2 affect MAD phosphorylation in the developing wing

To define the effects of the ISR on more proximal readouts of dpp signaling, we next examined MAD phosphorylation in pupal wings. During pupation, longitudinal veins are specified by epidermal growth factor receptor and dpp signaling (Blair, 2007). After the longitudinal veins have formed, the ACV and PCV are generated in response to Dpp that is transported from the adjacent longitudinal veins (Matsuda and Shimmi, 2012, Matsuda et al., 2013). As expected, 30 hours after pupariation, pMAD staining was detected in the presumptive ACV and PCV territories of driver control wings (Fig 2A, left panel). When dPPP1R15 was knocked down in the posterior compartment of the wing using *en>ppp1r15* RNAi, pMAD staining was evident in the PCV provein but was absent from ACV territory (Fig 2A, middle panel, ACV territory indicated by open triangle), whereas when dGCN2 was instead depleted in the posterior compartment using *en>gcn2* RNAi, ectopic pMAD staining was detected between the L4 and L5 proveins (Fig 2A, right panel, closed triangle). These changes in the distribution of MAD phosphorylation correlated well with the venation phenotypes observed in the adult wings of escapers (Fig 1A).

**Figure 2.**
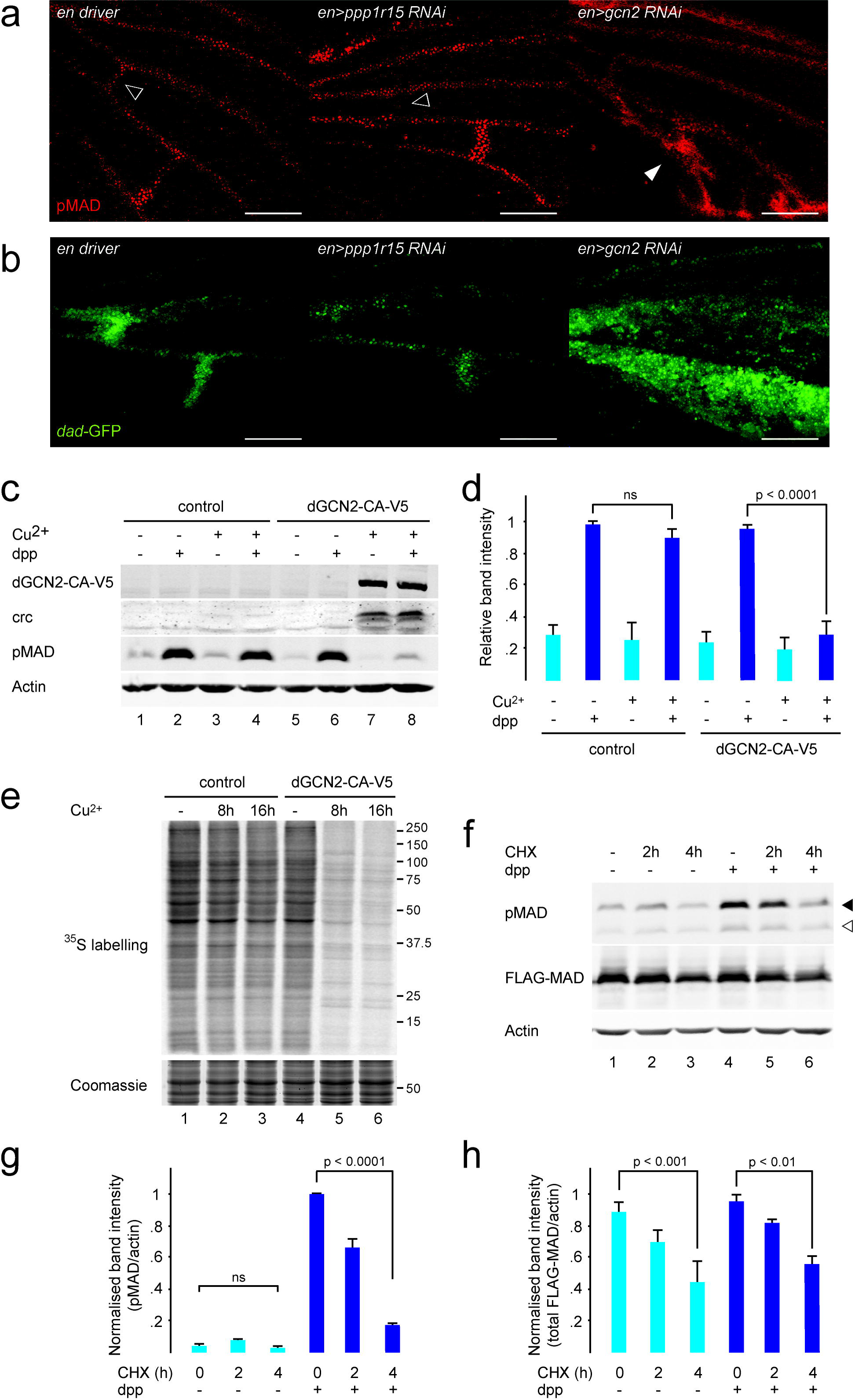
dPPP1R15 or dGCN2 affect MAD phosphorylation in the developing wing. (A) Representative fluorescence micrographs of pupal wings of the indicated genotypes at 30 hours after pupariation stained red for pMAD. Open arrowheads indicate ACV territory. Closed arrowheads indicate ectopic pMAD signal. Scale bars = 100µm. (B) Representative fluorescence micrograph of pupal wings of the indicated genotypes at 30 hours after pupariation. Green fluorescence indicates activation of the *dad*-*GFP.N* reporter. Scale bars = 100µm. (C) Immunoblot of cell lysates: lanes 1-4 S2 cells stably transfected with V5.pMT-Puro; lanes 5-8 S2 cells stably transfected with dGCN2-CAV5.pMT-Puro. Cu^2+^ indicates treatment with 0.7 mM copper sulphate for 16 hours. Dpp indicates treatment 1 nM Dpp for 1 hour prior to lysis. dGCN2-CA-V5 was detected with anti-V5 antibody. crc, pMAD and actin were detected using specific antibodies. (D) Quantification of pMAD staining in (C) with strongest signal with each experiment set as 1. n = 3. P value calculated using analysis of variance (ANOVA) with Bonferroni *post hoc* testing. (E) S2 cell lysates: lanes 1-3 of V5.pMT-Puro S2 cells; lanes 4-6 of dGCN2-CA-V5.pMT-Puro S2 cells. Cu^2+^ indicates treatment with 0.7 mM copper sulphate for the indicated times. ^35^S-labelled cysteine and methionine were added to cells for 10 minutes prior to lysis. “^35^S-labelled” indicates autoradiograph. “Coomassie” staining served as a loading control. (F) Immunoblot of S2 cell lysates expressing FLAG-MAD. CHX indicates treatment with 14 µg/ml cycloheximide for the indicated times. Dpp indicates treatment 0.5 nM nM dpp for 1 hour prior to lysis. FLAG-MAD was detected with an anti-FLAG antibody. pMAD and actin were detected with specific antibodies. Filled arrowheads indicate phosphorylated MAD-FLAG; open arrowheads indicate endogenous pMAD. (G) Quantification of phosphorylated FLAG-MAD (pMAD) and (H) total FLAG-MAD from (F), both normalised to actin signal with the strongest signal in each experiment set as 1. n = 3. P value calculated using ANOVA with Bonferroni *post hoc* testing.

To determine if the altered distribution of pMAD had functional consequences, we used a reporter comprising the promoter of a dpp-sensitive gene, *dad*, fused to the coding sequence of *GFP* (Hamaratoglu et al., 2011). As expected, in driver controls the GFP reporter signal was detected at the regions of the ACV and PCV proveins 30 hours after pupariation (Fig 2B, left panel). When *ppp1r15* was knocked down by *en>ppp1r15* RNAi, GFP signal was undetectable in the ACV provein territory (Fig 2B, middle panel), but when dGCN2 was depleted with *en>gcn2* RNAi, widespread ectopic reporter activation was seen, especially in the L4-L5 intervein region, and there was broadening of GFP signal into the L3-L4 intervein region (Fig 2B, right panel). Together, these data show that the precise arrangement of dpp signaling required for normal vein distribution in the pupal wing is dependent upon an intact ISR.

Because our *in vivo* studies had suggested that MAD phosphorylation is inhibited by activation of the ISR, we next turned to an *in vitro* model of dpp signaling to determine the mechanism of this interaction. Schneider-2 (S2) cells were generated to conditionally express a constitutively active dGCN2 tagged with the V5 epitope, dGCN2-CA-V5. In the absence of dGCN2-CA-V5, treatment with Dpp caused robust phosphorylation of MAD (Fig 2C & 2D). Induction of dGCN2-CA-V5 for 16 hours was sufficient to activate the ISR as evidenced by expression of the transcription factor crc (dATF4). Remarkably, expression of dGCN2-CA-V5 abolished dpp-induced phosphorylation of MAD (Fig 2C lanes 7 & 8, Fig 2D).

Activation of the ISR inhibits translation initiation (Dalton et al., 2012). Metabolic labeling with [^35^S]-methionine and cysteine confirmed that expression of dGCN2-CA-V5 for 8 or 16 hours reduced global translation (Fig 2E). It seemed plausible that loss of total MAD protein might therefore contribute to the loss of pMAD following induction of dGCN2-CA-V5. There are no antibodies that detect total MAD, so to estimate its half-life we transfected S2 cells with FLAG-tagged MAD and inhibited protein synthesis with cycloheximide (Fig 2F-H). Consistently, the level of total FLAG-MAD had halved by 4 hours after inhibiting translation but was insensitive to dpp. (Fig 2F & 2H). The levels of pMAD and phosphorylated FLAG-MAD were much lower than half of their starting level by 4 hours after treatment with cycloheximide (Fig 2F & 2G). These results indicate that activation of dGCN2 is sufficient to inhibit global protein synthesis and that inhibition of translation is sufficient to decrease the levels of both MAD and pMAD. The apparently preferential effect of translational attenuation on the levels of pMAD suggested, however, that additional short-lived proteins may be required for efficient MAD phosphorylation or that pMAD is preferentially destabilised.

### Crc regulates wing venation and antagonises MAD phosphorylation

crc is a bZIP transcription factor sharing sequence and functional homology with mammalian ATF4 (Hewes et al., 2000, Kang et al., 2015). In order to confirm activation of the ISR, we generated an antibody able to detect endogenous crc by western blot (Fig 2C and Supplementary Fig S2). This recognised a doublet of 65-70 kDa. After *in vitro* treatment with lambda phosphatase, crc doublets collapsed to a single band indicating that, like ATF4, crc is a phosphoprotein (Supplementary Fig S2). Similar to ATF4, the 5’ untranslated region (5’UTR) of the *crc* mRNA contains several small upstream open reading frames (uORFs), the last of which overlaps out-of-frame with the crc coding sequence (Supplementary Fig S2B). To confirm the observation by Kang et al (2015) that translation of *crc* is regulated in a manner similar to ATF4, we generated a reporter construct comprising the 5’UTR of *crc* fused to the coding sequence of luciferase. The reporter or a control consisting of a luciferase coding sequence lacking the *crc* 5’UTR were expressed in mammalian HEK293T cells and the ISR was activated using tunicamycin (Supplementary Fig S2C). The translation of the *crc*-reporter luciferase mRNA rose upon treatment with tunicamycin, while translation of the control fell. The ISR mediates its inhibitory effects on global translation through phosphorylation of eIF2α, rendering it an inhibitor of its own guanine nucleotide exchange factor, eIF2B (Vazquez de Aldana et al., 1993). The inhibition of eIF2B is also ultimately responsible for the increased translation of ATF4. These effects can be overcome in mammalian cells by the eIF2B-activating drug ISRIB (Sekine et al., 2015, Sidrauski et al., 2015). We therefore treated the HEK293T cells with ISRIB and observed a selective reduction of the translation of the *crc*-luciferase reporter (Supplementary Fig S2C).

We had previously shown that overexpression of the ISR kinase dPERK in the eye imaginal disc *(gmr>perk)* impairs eye development (Malzer et al., 2010). To test if this effect of the ISR on development might be mediated by crc, we expressed *gmr>crc* RNAi simultaneously with *gmr>*perk (Supplementary Fig S2D). This rescued eye growth and confirmed crc as a mediator of the ISR in *Drosophila*.

The ACV was unaffected when crc was depleted in the developing wing using *en>crc* RNAi, but *en>crc* RNAi suppressed the ACV phenotype of *en>ppp1r15* RNAi (Fig 3A & 3B). In the presence of dicer2, RNAi against *crc* driven by *enGAL4* caused loss of the ACV’s posterior portion similar to that observed with depletion of *gcn2* (Supplementary Fig S2E & 2F). Similar results were obtained with a whole-wing *nab* driver (Supplementary Fig S2G). *In situ* hybridization was performed to examine the distribution of *crc* mRNA in the developing wing (Fig 3C & Supplementary Fig S2H). In wing imaginal discs, *crc* expression was widespread throughout the pouch (Supplementary Fig S2H), while the pupal wing showed staining along the wing margin and surrounding the presumptive longitudinal and cross-veins (Fig 3C). Similar results were obtained using a second probe targeting a separate region of the *crc* mRNA (not shown).

**Figure 3.**
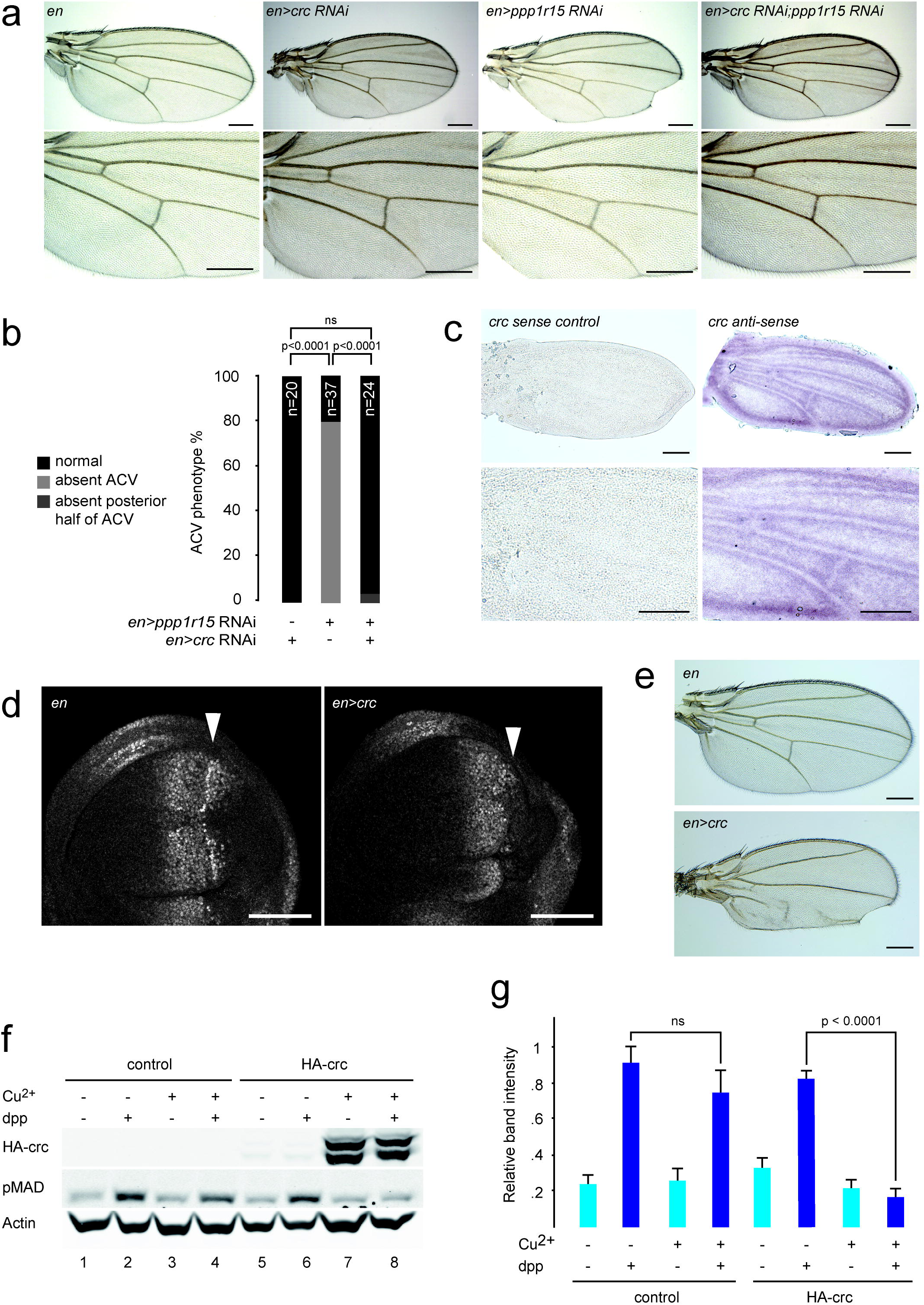
crc regulates wing venation and antagonises MAD phosphorylation. (A) Representative photomicrographs (5x objective) of adult wings of the indicated genotypes. *En* indicates *enGAL4* driver control. *en>crc* RNAi indicates *enGAL4>UAS-crc* RNAi. *en>ppp1r15* RNAi indicates *enGAL4>UAS-ppp1r15* RNAi. *en>ppp1r15* RNAi *;crc* RNAi indicates *enGAL4>UAS-crc RNAi;UAS-ppp1r15* RNAi. Lower panels are enlargements of the crossvein territories. Scale bars = 250µm. (B) Quantification of ACV phenotype in (A). P values calculated using ^2^ statistics with Bonferroni correction for multiple comparisons. (C) *In situ* hybridisation of w^1118^ pupal wings with sense or antisense probes to residues 1405-1900 of *crc* transcript A. Scale bars = 250µm. (D) Representative fluorescence micrograph (40x objective) of wing imaginal discs: signal = pMAD. *En* indicates *enGAL4* driver control. *en>crc* indicates *enGAL4>UAS-HA-crcA*. Orientation: left = anterior. Arrowhead indicates expected position of posterior pMAD zone. Scale bars = 50µm. (E) Representative photomicrographs of adult wings of the indicated genotypes. *En* indicates *enGAL4* driver control. *en>crc* indicates *enGAL4>UAS-crc*. Lower panels are enlargements of the crossvein territories. Scale bars = 250µm. (F) Immunoblot of S2 cell lysates: lanes 1-4 S2 cells stably transfected with HA.pMT-Puro; lanes 5-6 S2 cells stably transfected with HA-crcA.pMT-Puro. Cu^2+^ indicates treatment with 0.7 mM copper sulphate for 24 hours. dpp indicates treatment 0.5 nM dpp for 1 hour prior to lysis. HA-crc was detected with anti-HA antibody. pMAD and actin were detected using specific antibodies. (G) Quantification of pMAD staining in (F) with highest signal per experiment set as 1. n = 4. P value calculated using analysis of variance (ANOVA) with Bonferroni *post hoc* testing.

Next, we generated transgenic flies overexpressing crc. Wing imaginal discs expressing crc in the posterior compartment using the *enGAL4* driver showed reduced tissue mass and an absence of pMAD in the posterior portion of the disc (Fig 3D). In adult wings, crc expression in the posterior compartment of the wing reduced blade size and impaired venation (Fig 3E). When expressed in the whole wing using *nabGAL4*, crc generated smaller wings with evidence of inadequate crossvein, L3, L4 and L5 formation (Supplementary Fig S2I). These results indicated that crc can modify signals regulating venation *in vivo*. To examine this further, we generated S2 cells that conditionally expressed crc. As we had seen for dGCN2, crc expression blocked the phosphorylation of MAD caused by dpp (Fig 3F & 3G).

These results suggest that crc mediates at least some of the inhibition of BMP signalling that is caused by eIF2α hyperphosphorylation, and that crc is capable of attenuating MAD phosphorylation.

### 4E-BP mediates part of the crc effect on wing venation and MAD phosphorylation

To characterize the genes whose expression was altered by crc, we performed transcriptional profiling of S2 cells expressing crc for 3 or 6 hours (Fig 4A, & Supplementary Table). As expected, pathway analysis showed crc to induce genes involved in amino acid sufficiency and ribosome function (Supplementary Fig S3 & Supplementary Table). Gene Ontology term enrichment revealed the induction of many additional factors affecting translation (Supplementary Table). Transcripts that were significantly reduced included positive regulators of the cell cycle and nucleic acid biogenesis. Similar transcriptional changes were induced by expression of dGCN2-CA-V5 (Supplementary Fig S3; Supplementary Table). In contrast to dGCN2-CA-V5, an inactive mutant of dGCN2 (dGCN2-K552RV5) failed to induce genes involved in ribosome biogenesis, suggesting that increased protein synthetic load was not responsible for these effects (not shown).

**Figure 4.**
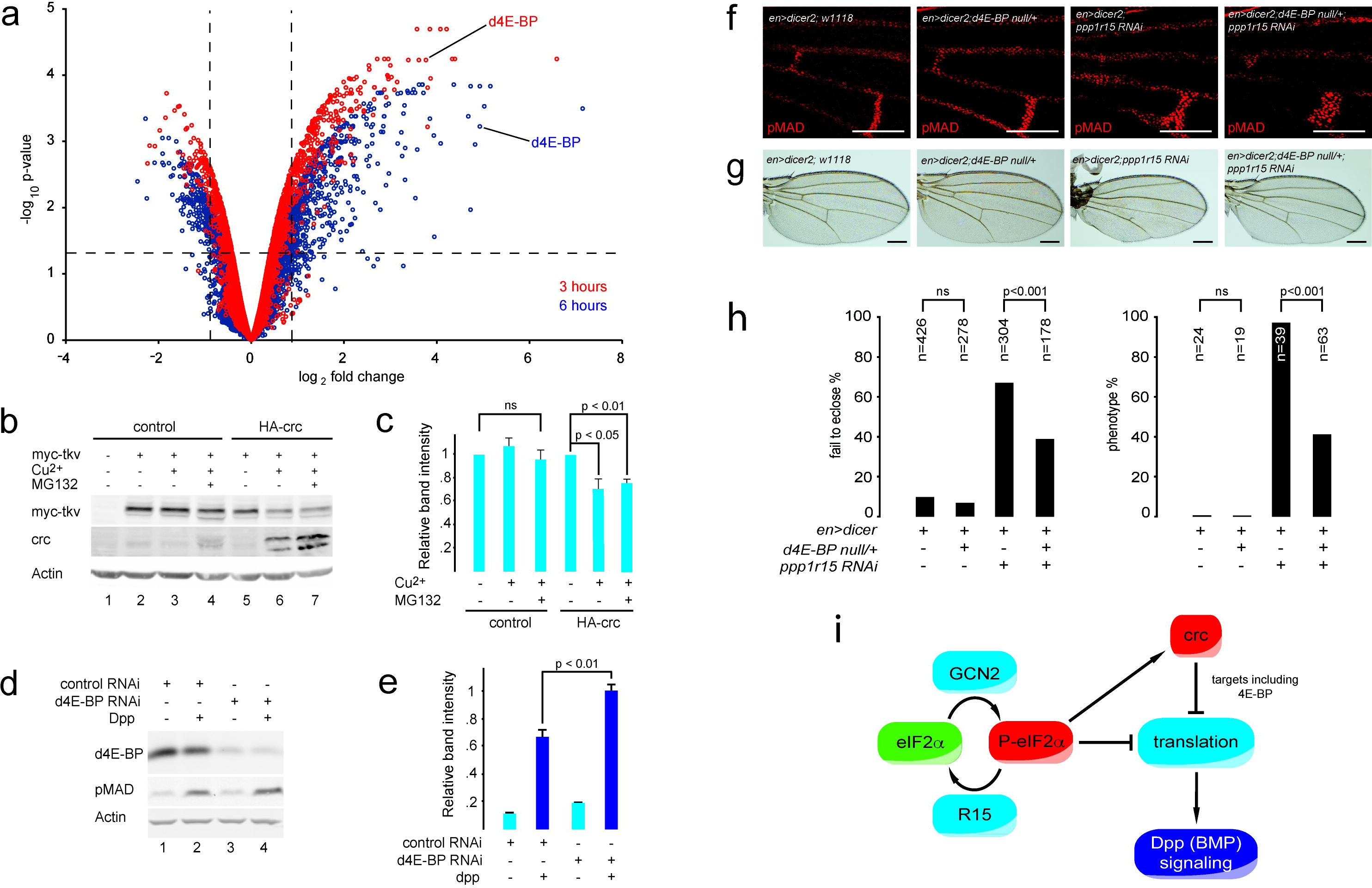
4E-BP contributes to the inhibition of MAD phosphorylation. (A) Microarray analysis of transcriptional changes caused by expression of crc in S2 cells. Volcano plot of transcriptional profiles of HA-crcA.pMT-Puro S2 stable cells relative to HA.pMT-Puro S2 stable cells, each treated with 0.7 mM copper sulphate for 3 hours (red symbols) or 6 hours (blue symbols). Vertical broken lines indicate 2^-/+0.7^-fold change. Horizontal broken line indicates p = 0.05 threshold. d4E-BP is indicated at 3 hours (red) and 6 hours (blue). (B) Immunoblot of cell lysates expressing myc-Tkv in the absence or presence of crc. (C) Quantification of (B) n = 3. P value calculated using ANOVA with Bonferroni *post hoc* testing. (D) Immunoblot of S2 cell lysates to assess the effect of d4E-BP small interfering RNA (RNAi) on MAD phosphorylation caused by 0.5 nM Dpp concentrations. (E) Quantification of (D) n = 3. P value calculated using ANOVA with Bonferroni *post hoc* testing. (F) Representative fluorescence micrographs of pupal wings of the indicated genotypes at 30 hours after pupariation stained red for pMAD. Scale bars = 100µm. (G) Representative photomicrographs (5x objective) of adult wings of the indicated genotypes. Scale bars = 200µm. (H) Quantification of animals from (G). Left graph indicates proportion of animals failing to eclose by 14. Right graph indicates frequency of wing vein phenotype if eclosing adults. P values calculated using ^2^ statistics with Bonferroni correction for multiple comparisons. (I) Schematic of interaction between integrated stress response (ISR) and BMP signaling. eIF2α is phosphorylated by GCN2 to P-eIF2α; PPP1R15 (R15) dephosphorylates P-eIF2α. P-eIF2α directly inhibits most cap-dependent translation of mRNAs, but induces expression of *crc* (*Drosophila* ATF4). Targets of crc further affect translation, e.g. 4E-BP antagonises translation of some mRNAs. On-going translation is necessary for efficient BMP signaling and so repression of protein synthesis by the ISR inhibits BMP signaling.

The preponderance of regulators of translation among the crc-sensitive transcripts raised the possibility that MAD phosphorylation might be affected in crc-expressing cells through additional changes to protein synthesis over and above those caused by eIF2α phosphorylation. An antibody capable of detecting the endogenous type I BMP receptor Tkv is lacking, so to determine if crc-induced inhibition of translation might affect Tkv protein levels we expressed myc-tagged Tkv in the inducible crc-expressing S2 cells. crc significantly suppressed myc-Tkv protein levels by approximately 20% and this could not be rescued by inhibition of the proteasome with MG132, indicating an effect on synthesis rather than proteasomal degradation of the protein (Fig 4B & 4C).

Although it is likely that many crc-sensitive factors cooperate to achieve this effect on protein synthesis, we chose to focus on eIF4E-binding protein (4E-BP) as it was one of the most highly induced negative regulators of translation in our transcriptional profiling (Fig 4A & Supplementary Table). The *Drosophila* homologue of 4E-BP (Thor) was up-regulated 30-fold at the mRNA level after 6 hours of crc expression (Fig 4A & Supplementary Fig S3D). This induction was confirmed at the protein level by western blot (Supplementary Fig S3E). Depletion of d4E-BP by RNAi in S2 cells significantly augmented dpp-induced MAD phosphorylation suggesting that d4E-BP exerts a tonic inhibition on dpp-MAD signaling (Fig 4D & 4E).

To test the relevance of this effect *in vivo*, we generated animals haploinsufficient for *d4E-BP*. In *d4E-PB^null/+^*(Tettweiler et al., 2005), the phosphorylation of MAD within the pupal wing vein territories was normal, as were the adult wing veins (Fig 4F and 4G). However, loss of one *d4E-BP* allele significantly rescued both the numbers of animals eclosing and the normal formation of the ACV in wings depleted of *ppp1r15* in the posterior compartment using *en>ppp1r15* RNAi (Fig 4F-4H).

These findings indicate that the impaired dpp-MAD signaling observed in this model is sensitive to the levels of d4E-BP. Taken together, our observations suggest that targets of crc that regulate translation contribute to the inhibition of dpp signaling during development.

## Discussion

We have shown that the ISR modulates tissue morphogenesis through the regulation of dpp-induced MAD phosphorylation. In wing tissue, this mechanism is driven primarily by the eIF2α kinase dGCN2. These repressive effects are achieved directly by the reduction in translation that accompanies phosphorylation of eIF2α, and indirectly by the induction of the transcription factor crc (dATF4) and its targets including d4E-BP (Fig 4I). Since the ISR is conserved between metazoans, our findings may have wider significance in developmental biology. Moreover, because mutations of the BMP receptor BMPR2 and the stress sensing kinase GCN2 account for the majority of inherited cases of human PAH, this interaction may have relevance for the development of new therapies for this currently incurable disease.

Developmental signals orchestrate tissue patterning by following predetermined programmes. Environmental factors also impact on development and so crosstalk between stress signaling and developmental pathways is necessary. It is known that overexpression of non-phosphorylatable mutants of eIF2α accelerates development of enlarged adult female flies, while expression of a phosphomimetic eIF2α delays larval development (Qu et al., 1997). We previously reported that depletion of the eIF2α phosphatase dPPP1R15 causes a developmental delay similar to that of phosphomimetic eIF2α (Malzer et al., 2010). We have now shown that expression of dPPP1R15 is necessary for larval development only in specific larval tissues, including the imaginal discs, and shares an antagonistic relationship with dGCN2.

*In vitro* studies indicate that inhibition of protein synthesis mediates some of the inhibitory effects of dGCN2 on BMP signaling, reflecting the short half-lives of components of the BMP signaling cascade. Vein formation in the fly wing is governed by BMP signaling. dpp (the *Drosophila* BMP2/4 homologue) binds to the type I receptors, Tkv or Sax, and type II receptor Punt to phosphorylate and activate the transcription factor MAD (Arora et al., 1995). Crossvein morphogenesis requires secretion of dpp from nearby longitudinal veins and its chaperoning by the molecules tsg, cv and sog, which are subsequently degraded by Tlr to release dpp at sites defined by high levels of cv-2 (Shimmi et al., 2005, Matsuda and Shimmi, 2012). The formation of the dpp gradient also requires the expression of extracellular glypicans, such as dally, and their post-translational modification by enzymes including sulfateless (Vuilleumier et al., 2010, Kirkpatrick et al., 2006). Changes in the expression levels of at least some of these components may contribute to impaired BMP signaling during activation of the ISR. *dally* RNAi had a more dramatic effect on wing development when expressed with *ppp1r15* RNAi, compared with *ppp1r15* RNAi in flies with one hypomorphic allele of *dpp^d5^*. This might relate to differences in the degree to which *dally* and *dpp* were depleted, but might also reflect the dual role of dally in both stabilizing and dispersing dpp in the extracellular space and as a co-receptor involved directly in dpp signaling (Hamaratoglu et al., 2014). Examination of wing imaginal discs has not yet revealed dramatic effects of the ISR on signaling via the wnt or hedgehog pathways (not shown), but further studies are necessary before the regulation of developmental signaling by the ISR can be said to show specificity towards the BMP pathway.

crc, the *Drosophila* homologue of ATF4, also inhibits MAD phosphorylation. The large number of genes sensitive to crc suggests that its effect on BMP signaling may be multifaceted. Our data reveal, however, that part of this effect is mediated by the induction of d4E-BP. Of note, ATF4 binding sites have recently been identified within the *d4E-BP* gene (Kang et al., 2017). By binding to eIF4E, 4E-BP prevents assembly of eIF4F and so selectively inhibits cap-dependent translation (Sonenberg and Hinnebusch, 2009). Interestingly, expression of a hyperactive mutant of d4E-BP in the wing has been shown to result in selective loss of the ACV, although the mechanism was unknown (Miron et al., 2001). How elevated d4E-BP levels inhibit phosphorylation of MAD in the absence of detectable effects on global translation rates is unclear. It is plausible that the extent of translational attenuation may vary among cap-dependent mRNAs and, in such a model, as levels of available eIF4E declined some mRNAs might compete more efficiently than others for a limited supply of the eIF4F. Such sensitivity could explain the some of the effects we have described, although the mRNAs responsible for altered MAD phosphorylation have yet to be fully identified. Nevertheless, there are numerous instances in which d4E-BP selectively regulates mRNA translation. For example, insulin signaling inhibits neurotransmitter release via d4E-BP mediated repression of *complexin* mRNA translation (Mahoney et al., 2016), while dietary restriction enhances the expression of mitochondrial respiratory components by inducing d4E-BP (Zid et al., 2009). Indeed, there is emerging evidence in *Drosophila* that ISR-induced d4E-BP plays a role in biasing translation during infection (Vasudevan et al., 2017), development and aging (Kang et al., 2017).

Mice generated to be insensitive to the ISR kinases owing to mutation of the target serine 51 of eIF2α revealed a role for the ISR in mammalian development (Scheuner et al., 2001). Homozygous pups were growth retarded and died from hypoglycemia due to impaired gluconeogenesis, while heterozygous animals developed diabetes if fed high fat chow owing to impaired pancreatic ß-cell survival. Mutations of GCN2 in humans have now been implicated in the pathogenesis of PAH, a disease known to involve aberrant BMP signaling that permits excessive proliferation of pulmonary arterial smooth muscle cells (Eyries et al., 2014, Best et al., 2014). Why loss of GCN2-mediated inhibition of BMP signaling should cause a disorder more commonly associated with insufficient SMAD phosphorylation is intriguing. However, mammalian BMP signaling is more complex than that of insects and it is known that loss of signaling via one BMP type 2 receptor in pulmonary artery smooth muscle cells can lead to excessive signaling through other type 2 receptors (Yu et al., 2005).

In summary, we report a novel mechanism for the modulation of BMP signaling by the ISR. This involves direct modulation of translation initiation through eIF2α phosphorylation and indirect effects via the crc-d4E-BP axis. This raises the possibility that pharmacological manipulation of the ISR may represent a therapeutic approach for the regulation of BMP signaling.

## Materials and Methods

### Drosophila genetics

The following strains were obtain from the Vienna Drosophila RNAi Center: *ppp1r15* (RNAi #1:15238; RNAi #2: v107545); *gcn2* (v103976); *crc* (v109014); *dally* (14136) and 51D background as a control line. Stocks obtained from the Bloomington Drosophila Stock Center (NIH P40OD018537) were: *UAS-dally* (5397); *engrailed-Gal4* (6356); *UAS-dicer2*; *en-Gal4, UAS-eGFP* (25752); *UAS-dpp* (1486); *dpp^d5^* (2071); *dpp^hr56^* (36528); *GMR-Gal4* (1104). Other lines were supplied by: isogenic w^1118^ line; w^1118^; *if/CyO*; *gmr-GAL4/TM6B* (Dr S Imarisio; University of Cambridge); *escargot^NP7397^-Gal4; yw.hs-flp122*; *Act5c>y^+^>Gal4, UAS-GFP*; *MKRS/TM6b, tb* (Dr J de Navascues Melero; University of Cardiff); nab^NP3537^-Gal4 (Prof S Russell; University of Cambridge); UAS-dGcn2-CA (Dr P Leopold, University of Nice) (Bjordal et al., 2014); *d4E-BP^Null^* line (Dr J Carmichael; University of Cambridge) (Tettweiler et al., 2005); *dad-GFP.N* (Ninov et al., 2010); the *uas-perk* line was described previously (Malzer et al., 2010).

Unless stated otherwise, crosses were performed at 25°C with three to four virgins and two males in standard food vials. Each two to four days these flies were then flipped into fresh vials to avoid overcrowding of progeny. The food used was a standard ‘lower maize, higher yeast agar’ recipe consisting of 2% (w/v) yeast, 8% (w/v) dextrose, 7% (w/v) maize and 1% (w/v) agar with the addition of nipagin and dry yeast pellets. In specific experiments modified foods were used: “high protein food” [5.9% (w/v) glucose, 6.6% (w/v) cornmeal, 4% (w/v) dried yeast and 0.7% agar] or “low protein foods” [5.9% (w/v) glucose, 6.6% (w/v) cornmeal, 0.25% (w/v) dried yeast and 0.7% agar].

For the tissue-specific screen, virgin females of the *ppp1r15* RNAi #1 or w^1118^ were crossed to males of various GAL4-driver lines. Fourteen days after egg laying (AEL), progeny were analysed. Developmental was performed as described previously (Malzer et al., 2013). To generate flip out clones in wing imaginal discs, we crossed yw.hs-flp^122^; *act5c>y^+^>Gal4, UAS-GFP; MKRS/TM6b, tb to either w1118 (control), ppp1r15 RNAi* #1 or *UAS-dGcn2-CA* flies. Vials were heat shocked 4 days AEL for 15 minutes at 37°C. The following day, wing imaginal discs of non-tubby third instar larvae were dissected.

### Immunohistochemistry

Larval wing imaginal discs were dissected in PBS and fixed with 4% paraformaldehyde in PBS for 30 minutes at room temperature, followed by washes with PBT (PBS, 0.1% Triton X-100). For pupal wing dissections, pupae were collected at the appropriate number of hours after puparium formation (APF) and fixed with an opened case over night at 4°C with 4% paraformaldehyde in PBS. After dissection, an additional fixation for 30 minutes at room temperature was performed. Tissues were stained with the primary rabbit anti-pSMAD antibody (PS1) 1:500 (from Prof P. ten Dijke, University of Leiden) overnight at 4°C followed by anti-rabbit Alexa 594 1:250/500 (Thermo Fisher Scientific) for 1 hour at room temperature. Samples were mounted in ProLong Gold Antifade with DAPI (Thermo Fisher Scientific). Images were taken using a Zeiss LSM880 microscope with a 20x and 40x objective. Merged images of Z-stack focal planes were generated with ImageJ (NIH) showing maximum intensity.

### Generation of transgenic flies

The *UAS-HA-crcA* line was generated by amplification of the *HA-crcA* sequence from the construct HA-crcA.pMT-Puro and directionally cloned between NotI and XhoI into pUASTattB. Microinjection was performed by Department of Genetics core facility, University of Cambridge, and stock number 13-14 yielded an insertion on the third chromosome (86F8).

### Expression plasmids

The HA-tag sequence was directionally cloned between BamHI and EcoRI into pcDNA3.1 (HA.pcDNA3.1) and then sub-cloned between KpnI and XhoI of pMT-Puro vector (Addgene 17923) to generated HA.pMT-Puro. Crc transcript A coding sequence was amplified from cDNA clone RH01327 (DGRC, Indiana University, USA) and directionally cloned between EcoRI and XhoI into HA.pcDNA3.1 plasmid then HA-crcA was sub-cloned between KpnI and XhoI into pMT-Puro vector (Addgene 17923) to generate HA-crcA.pMT-Puro. To generate dGCN2-CA-V5.pMT-Puro, the *gcn2* coding sequence was amplified from the cDNA clone AT10027 (DGRC, Indiana University, USA) and mutated to incorporate an activating mutation in the translated protein (F751L) then cloned into pMT-Puro vector (from David Sabatini, Addgene stock 17923). To generate the 5’UTR-*crcELuciferase* reporter-construct, a synthesised DNA-fragment (GeneArt, Thermo Fisher) containing the 5’UTR of *crcE* and the first three amino acids of the protein coding sequence was cloned in frame into a Luciferase-pcDNA3.1 plasmid (Malzer et al., 2013) by Gibson assembly. The crc-pGEX-6P-1 expression construct was generated by amplifying the crcA coding sequence from the cDNA clone RH01327 (DGRC, Indiana University, USA) following by cloning between SalI and NotI in pGEX-6P-1 (Invitrogen). The construct pAFW-MAD-FLAG (Kunnapuu et al., 2009) was used to express MAD-FLAG; the construct myc-tkv.pAc5.1 was used to express myc-tkv and was generated from the myc-tkv.pMT plasmid (Li et al., 2016). For punt-V5 expression, the punt coding sequence was amplified from plasmid FMO13005 (DGRC, Indiana University, USA) and cloned between KpnI and XhoI into pAc5.1 plasmid (Thermo Fisher); for myc-sax expression, sax coding sequence was amplified from plasmid 02439 (DGRC, Indiana University, USA) and similarly cloned in pAc5.1.

### S2 cell culture

Cycloheximide was from Sigma Aldrich; dpp was R&D Systems. *Drosophila* Schneider S2 cells (from Dr J Hirst, Cambridge) were grown at 25°C in Schneider medium (Sigma-Aldrich) supplemented with 10% fetal bovine serum (Invitrogen) and 100 U/ml streptomycin/penicillin (Sigma-Aldrich). Transfection reagent TransIT 2020 (Mirus Bio) was used for all experiments. To generate stable inducible lines, S2 cells were transfected with dGCN2-CA-V5.pMT-Puro or HA-crcA.pMT-Puro constructs and cultured for 2 weeks in 4 µg/ml puromycin. In parallel, control cell lines were generated with pMT-Puro or HA.pMT-Puro. Transgene expression was induced with 0.7 mM copper sulphate. For measurement of Dpp signaling, 2.5×10^6^ cells per well were seeded in 6-well plates and expression was induced for 16 hours (dGCN2-CA-V5) or 24 hours (HA-crcA), followed by treatment with 0.5 nM or 1 nM Dpp for 1 hour. When assessing protein half-lives, S2 cells were transfected in 6-well plates with 250 ng of myc-tkv.pAC5.1, myc-sax pAC5.1, or punt-V5.pAC5.1. Twenty-four hours after transfection, cells were treated with 100 µg/ml cycloheximide for up to 12 hours as indicated. To assess level of pMAD-FLAG and total MAD-FLAG, S2 cells were transfected with 1 µg of MADFLAG.pAFW. Twenty-four hours post-transfection, cycloheximide (14 µg/ml or 100 µg/ml as indicated) was added for the indicated times with 1 nM dpp present for the final hour.

### Microarray

dGCN2-CA-V5 pMT-puro or HA-crc pMT-puro inducible cell lines induced with 0.7 mM copper sulphate for indicated times. pMT-puro and HA-pMT-puro lines were induced with 0.7 mM copper sulphate for control purposes. Total RNA was prepared from cells by homogenization and extraction using TRIzol reagent (GibcoBRL). Each total RNA sample (50 μg) was subjected to reverse transcription and direct labelling with Cy3- or Cy5-dCTPs (Amersham). Appropriate Cy3-dCTP or Cy5-dCTP labelled samples were mixed together and hybridised to the *Drosophila* Array Consortium (INDAC) oligo array FL003 for 16 hours at 51°C (Genetics core facility, University of Cambridge, UK). After hybridisation, slides were washed, spun dry and scanned with 635-nm and 532-nm lasers using a Genepix 4000B scanner (Axon Instruments). Spot intensities were normalised using variance stabilisation (Huber et al., 2002) in the Vsn package in R/Bioconductor. The magnitude and significance of each spot intensity was estimated using linear models in the LIMMA package in R/Bioconductor. False discovery rates (FDR) were calculated using the Benjamini-Hochberg method (Hochberg and Benjamini, 1990). Differentially expressed genes (exhibiting log_2_-fold changes <-0.7 or >0.7 and an FDR-adjusted p-value of <⁊0.05) were subjected to GO and KEGG pathway enrichment analysis using FlyMine (Lyne et al., 2007).

### Immunoblotting

S2 cells were lysed RIPA buffer (50 mM Tris-HCl pH7.4; 150 mM NaCl; 1% NP-40; 0.5% Sodium-deoxycholate; 0.1% SDS; 2mM EDTA) supplemented with 1mM PMSF and EDTA-free protease inhibitors (Sigma Aldrich). Commercially available primary antibodies used were: rabbit anti-phospho-SMAD 1/5 (which recognises *Drosophila* pMAD; 9516; Cell signaling); rabbit anti-Actin (A2066; Sigma-Aldrich); rabbit 4E-BP (4923; Cell signaling).

### Crc antibody preparation

BL21(DE3) pLysS *E. coli* were transformed with crc-pGEX-6P-1 then treated overnight at 37°C with 1 mM IPTG to induce expression. Recombinant protein was purified on Glutathione Sepharose 4B resin and eluted with PreScission protease (GE Healthcare). Rabbit polyclonal antibodies were generated by Cambridge Research Biochemicals, UK, using this antigen.

### *In situ* hybridisation

The 3’UTR of *crcA* was amplified (residues 1405-1900) from the *crcA* cDNA clone RH01327 (DGRC, Indiana University, USA) and cloned into pcDNA3 (Invitrogen) by Gibson Assembly (New England Biolabs). Antisense and sense DIG-labeled RNA probes were synthesised from linearised plasmid DNA using a SP6/T7 DIG-RNA labelling kit (Roche Molecular Biochemicals, Mannheim, Germany). Wing imaginal discs and pupal wings were dissected in PBS and fixed in 4% paraformaldehyde in PBS for 20 minutes at room temperature, washed twice with PBT and once with methanol. Fixed samples were washed twice with ethanol and incubated in a mixture of xylene and ethanol (1:1 v/v) for 60 minutes, washed twice in ethanol and rehydrated by immersion in a graded methanol series (80%, 50%, 25% v/v in water) and then water. Samples were treated with acetone (80%) at −20°C then washed twice with PBT. They were fixed again in 4% paraformaldehyde before being washed further with PBT then incubated at room temperature with 1:1 PBT:hybridization buffer (HB, 50% formamide, 5X SSC, 5X Denharts solution, 0.1% Tween 20, 100 µg/ml yeast tRNA, RNAse free water). They were pre-hybridised for 3 hours in HB at 60°C. Sense and antisense riboprobes were diluted 1:1000 in HB and denatured at 80°C. Samples were hybridised with diluted riboprobes at 60°C for 18 hours. The following day, samples were washed with HB solution at 60°C before then sequentially in 50% and 25% HB solution (v/v) in PBT. Following further washes in PBT, the hybridised probes were detected using anti-DIG-alkaline phosphatase conjugated sheep IgG (Fab fragments) secondary antibody using NBT/BCIP chromogenic substrates (Roche Molecular Biochemicals, Mannheim, Germany).

### ^35^S labelling of cultured S2 cells

Expression of dGCN2-CA-V5 or HA-crc was induced in stable S2 cell lines by treatment with 0.7mM copper sulphate. Thirty minutes prior to cell harvest, ten million cells were washed in PBS and resuspended in 1ml of cysteine and methionine free Dulbecco Modified Eagle Medium (DMEM) (MP Biomedicals, US. Cat.1642454) supplemented with 10% dialysed FBS and 10% Schneider medium. ^35^S-labelled cysteine and methionine Easy Tag Express protein labelling mix (Perkin Elmer) were added to cells for the final 10 minutes of the time course before addition of 20 µg/ml cycloheximide and incubation on ice. Cells were harvested and washed in cold PBS containing 20µg/ml CHX, then lysed in harvest buffer (HEPES pH 7.9, 10 mM; NaCl 50 mM; sucrose 0.5M; EDTA 0.1 mM; 0.5% v/v Triton X-100) supplemented with protease inhibitor cocktail (Roche, Welwyn Garden City, UK) and 1 mM PMSF. Post nuclear supernatants were separated by SDS-PAGE on 12.5% acrylamide gels and stained InstantBlue Coomassie stain (Expedeon, San Diego, CA, USA). ^35^S incorporation was analysed by exposure to a phosphostorage plate.

### Luciferase Assay

To analyse the regulatory function of the 5’ UTR of the *crcE* mRNA, HEK293T cells were transfected with Luc-pcDNA3.1 or 5’UTRcrcE-Luc.pcDNA3.1 constructs and TK-Renilla luciferase plasmid as a transfection control. Six hours post-transfection, cells were treated for 16 hours with tunicamycin (2.5 µg/ml) and/or ISRIB (45 ng/ml). Control cells were treated with the appropriate vehicle controls. A Dual-Glo® Luciferase reporter assay (Promega, UK) was subsequently carried out according to the manufacturer’s instructions to quantify the fold-induction of luciferase upon drug treatment. The ratio of firefly/*Renilla* luciferase luminescence was calculated and expressed as fold change compared to untreated samples.

## Acknowledgements

SJM was supported by the MRC. EM was supported by the MRC and Wellcome Trust ISSF. CSD was a Wellcome Trust PhD student. JEC was supported by the Alpha1 Foundation. The CIMR microscopy core facility is supported by a Wellcome Trust Strategic Award [100140] and a Wellcome Trust equipment grant [093026]. We thank David Ron for critical reading of the manuscript.

## References

Arora, K., Dai, H., Kazuko, S., Jamal, J., O’Connor, M., Letsou, A. & Warrior, R. 1995. The Drosophila *schnurri* gene acts in the Dpp/TGFb signalling pathway and encodes a transcription factor homologous to the human MBP family. Cell, 81, 781-790.

Belenkaya, T. Y., Han, C., Yan, D., Opoka, R. J., Khodoun, M., Liu, H. & Lin, X. 2004. Drosophila Dpp morphogen movement is independent of dynamin-mediated endocytosis but regulated by the glypican members of heparan sulfate proteoglycans. Cell, 119, 231-44.

Best, D. H., Sumner, K. L., Austin, E. D., Chung, W. K., Brown, L. M., Borczuk, A. C., Rosenzweig, E. B., Bayrak-Toydemir, P., Mao, R., Cahill, B. C., Tazelaar, H. D., Leslie, K. O., Hemnes, A. R., Robbins, I. M. & Elliott, C. G. 2014. EIF2AK4 mutations in pulmonary capillary hemangiomatosis. Chest, 145, 231-6.

Bjordal, M., Arquier, N., Kniazeff, J., Pin, J. P. & Leopold, P. 2014. Sensing of amino acids in a dopaminergic circuitry promotes rejection of an incomplete diet in Drosophila. Cell, 156, 510-21.

Blair, S. S. 2007. Wing vein patterning in Drosophila and the analysis of intercellular signaling. Annu Rev Cell Dev Biol, 23, 293-319.

Capdevila, J., Estrada, M. P., Sanchez-Herrero, E. & Guerrero, I. 1994. The Drosophila segment polarity gene patched interacts with decapentaplegic in wing development. EMBO J, 13, 71-82.

Dalton, L. E., Healey, E., Irving, J. & Marciniak, S. J. 2012. Phosphoproteins in stress-induced disease. Prog Mol Biol Transl Sci, 106, 189-221.

Delepine, M., Nicolino, M., Barrett, T., Golamaully, M., Lathrop, G. M. & Julier, C. 2000. EIF2AK3, encoding translation initiation factor 2-alpha kinase 3, is mutated in patients with Wolcott-Rallison syndrome. Nat Genet, 25, 406-9.

Eyries, M., Montani, D., Girerd, B., Perret, C., Leroy, A., Lonjou, C., Chelghoum, N., Coulet, F., Bonnet, D., Dorfmuller, P., Fadel, E., Sitbon, O., Simonneau, G., Tregouet, D. A., Humbert, M. & Soubrier, F. 2014. EIF2AK4 mutations cause pulmonary veno-occlusive disease, a recessive form of pulmonary hypertension. Nat Genet, 46, 65-9.

Hamaratoglu, F., Affolter, M. & Pyrowolakis, G. 2014. Dpp/BMP signaling in flies: from molecules to biology. Semin Cell Dev Biol, 32, 128-36.

Hamaratoglu, F., de Lachapelle, A. M., Pyrowolakis, G., Bergmann, S. & Affolter, M. 2011. Dpp signaling activity requires Pentagone to scale with tissue size in the growing Drosophila wing imaginal disc. PLoS Biol, 9, e1001182.

Harding, H., Zhang, Y., Zeng, H., Novoa, I., Lu, P., Calfon, M., Sadri, N., Yun, C., Popko, B., Paules, R., Stojdl, D., Bell, J., Hettmann, T., Leiden, J. & Ron, D. 2003. An integrated stress response regulates amino acid metabolism and resistance to oxidative stress. Mol Cell, 11, 619-633.

Harding, H. P., Zhang, Y., Scheuner, D., Chen, J. J., Kaufman, R. J. & Ron, D. 2009. Ppp1r15 gene knockout reveals an essential role for translation initiation factor 2 alpha (eIF2alpha) dephosphorylation in mammalian development. Proc Natl Acad Sci U S A, 106, 1832-7.

Hewes, R. S., Schaefer, A. M. & Taghert, P. H. 2000. The cryptocephal gene (ATF4) encodes multiple basic-leucine zipper proteins controlling molting and metamorphosis in Drosophila. Genetics, 155, 1711-23.

Hochberg, Y. & Benjamini, Y. 1990. More powerful procedures for multiple significance testing. Stat Med, 9, 811-8.

Huber, W., Von Heydebreck, A., Sultmann, H., Poustka, A. & Vingron, M. 2002. Variance stabilization applied to microarray data calibration and to the quantification of differential expression. Bioinformatics, 18 Suppl 1, S96-104.

International, P. P. H. C., Lane, K. B., Machado, R. D., Pauciulo, M. W., Thomson, J. R., Phillips, J. A., 3RD, Loyd, J. E., Nichols, W. C. & Trembath, R. C. 2000. Heterozygous germline mutations in BMPR2, encoding a TGF-beta receptor, cause familial primary pulmonary hypertension. Nat Genet, 26, 81-4.

Kang, K., Ryoo, H. D., Park, J. E., Yoon, J. H. & Kang, M. J. 2015. A Drosophila Reporter for the Translational Activation of ATF4 Marks Stressed Cells during Development. PLoS One, 10, e0126795.

Kang, M. J., Vasudevan, D., Kang, K., Kim, K., Park, J. E., Zhang, N., Zeng, X., Neubert, T. A., Marr, M. T., 2ND & Ryoo, H. D. 2017. 4E-BP is a target of the GCN2-ATF4 pathway during Drosophila development and aging. J Cell Biol, 216, 115-129.

Kirkpatrick, C. A., Knox, S. M., Staatz, W. D., Fox, B., Lercher, D. M. & Selleck, S. B. 2006. The function of a Drosophila glypican does not depend entirely on heparan sulfate modification. Dev Biol, 300, 570-82.

Kunnapuu, J., Bjorkgren, I. & Shimmi, O. 2009. The Drosophila DPP signal is produced by cleavage of its proprotein at evolutionary diversified furin-recognition sites. Proc Natl Acad Sci U S A, 106, 8501-6.

Li, J. & Li, W. X. 2006. A novel function of Drosophila eIF4A as a negative regulator of Dpp/BMP signalling that mediates SMAD degradation. Nat Cell Biol, 8, 1407-14.

Li, W., Yao, A., Zhi, H., Kaur, K., Zhu, Y. C., Jia, M., Zhao, H., Wang, Q., Jin, S., Zhao, G., Xiong, Z. Q. & Zhang, Y. Q. 2016. Angelman Syndrome Protein Ube3a Regulates Synaptic Growth and Endocytosis by Inhibiting BMP Signaling in Drosophila. PLoS Genet, 12, e1006062.

Lu, P. D., Harding, H. P. & Ron, D. 2004. Translation re-initiation at alternative open reading frames regulates gene expression in an integrated stress response. J Cell Biol, 167, 27-33.

Lyne, R., Smith, R., Rutherford, K., Wakeling, M., Varley, A., Guillier, F., Janssens, H., Ji, W., Mclaren, P., North, P., Rana, D., Riley, T., Sullivan, J., Watkins, X., Woodbridge, M., Lilley, K., Russell, S., Ashburner, M., Mizuguchi, K. & Micklem, G. 2007. FlyMine: an integrated database for Drosophila and Anopheles genomics. Genome Biol, 8, R129.

Ma, Y. & Hendershot, L. M. 2003. Delineation of a Negative Feedback Regulatory Loop That Controls Protein Translation during Endoplasmic Reticulum Stress. J Biol Chem, 278, 34864-73.

Machado, R. D., Pauciulo, M. W., Thomson, J. R., Lane, K. B., Morgan, N. V., Wheeler, L., Phillips, J. A., 3RD, Newman, J., Williams, D., Galie, N., Manes, A., Mcneil, K., Yacoub, M., Mikhail, G., Rogers, P., Corris, P., Humbert, M., Donnai, D., Martensson, G., Tranebjaerg, L., Loyd, J. E., Trembath, R. C. & Nichols, W. C. 2001. BMPR2 haploinsufficiency as the inherited molecular mechanism for primary pulmonary hypertension. Am J Hum Genet, 68, 92-102.

Mahoney, R. E., Azpurua, J. & Eaton, B. A. 2016. Insulin signaling controls neurotransmission via the 4eBP-dependent modification of the exocytotic machinery. Elife, 5.

Malzer, E., Daly, M. L., Moloney, A., Sendall, T. J., Thomas, S. E., Ryder, E., Ryoo, H. D., Crowther, D. C., Lomas, D. A. & Marciniak, S. J. 2010. Impaired tissue growth is mediated by checkpoint kinase 1 (CHK1) in the integrated stress response. J Cell Sci, 123, 2892-900.

Malzer, E., Szajewska-Skuta, M., Dalton, L. E., Thomas, S. E., Hu, N., Skaer, H., Lomas, D. A., Crowther, D. C. & Marciniak, S. J. 2013. Coordinate regulation of eIF2alpha phosphorylation by dPPP1R15 and dGCN2 is required during development. J Cell Sci, 126, 1406-15.

Marciniak, S. J., Yun, C. Y., Oyadomari, S., Novoa, I., Zhang, Y., Jungreis, R., Nagata, K., Harding, H. P. & Ron, D. 2004. CHOP induces death by promoting protein synthesis and oxidation in the stressed endoplasmic reticulum. Genes Dev, 18, 3066-3077.

Matsuda, S., Blanco, J. & Shimmi, O. 2013. A feed-forward loop coupling extracellular BMP transport and morphogenesis in Drosophila wing. PLoS Genet, 9, e1003403.

Matsuda, S. & Shimmi, O. 2012. Directional transport and active retention of Dpp/BMP create wing vein patterns in Drosophila. Dev Biol, 366, 153-62.

Miron, M., Verdu, J., Lachance, P. E., Birnbaum, M. J., Lasko, P. F. & Sonenberg, N. 2001. The translational inhibitor 4E-BP is an effector of PI(3)K/Akt signalling and cell growth in Drosophila. Nat Cell Biol, 3, 596-601.

Montani, D., Achouh, L., Dorfmuller, P., Le Pavec, J., Sztrymf, B., Tcherakian, C., Rabiller, A., Haque, R., Sitbon, O., Jais, X., Dartevelle, P., Maitre, S., Capron, F., Musset, D., Simonneau, G. & Humbert, M. 2008. Pulmonary veno-occlusive disease: clinical, functional, radiologic, and hemodynamic characteristics and outcome of 24 cases confirmed by histology. Medicine (Baltimore), 87, 220-33.

Ninov, N., Menezes-Cabral, S., Prat-Rojo, C., Manjon, C., Weiss, A., Pyrowolakis, G., Affolter, M. & Martin-Blanco, E. 2010. Dpp signaling directs cell motility and invasiveness during epithelial morphogenesis. Curr Biol, 20, 513-20.

Novoa, I., Zeng, H., Harding, H. & Ron, D. 2001. Feedback inhibition of the unfolded protein response by GADD34-mediated dephosphorylation of eIF2a. J Cell Biol, 153, 1011-1022.

Pereira, R. C., Stadmeyer, L., Marciniak, S. J., Ron, D. & Canalis, E. 2006. C/EBP homologous protein is necessary for normal osteoblastic function. Journal of cellular biochemistry, 97, 633-40.

Qu, S., Perlaky, S. E., Organ, E. L., Crawford, D. & Cavener, D. R. 1997. Mutations at the Ser50 residue of translation factor eIF-2alpha dominantly affect developmental rate, body weight, and viability of Drosophila melanogaster. Gene expression, 6, 349-60.

Ray, R. P. & Wharton, K. A. 2001. Context-dependent relationships between the BMPs gbb and dpp during development of the Drosophila wing imaginal disk. Development, 128, 3913-25.

Runo, J. R., Vnencak-Jones, C. L., Prince, M., Loyd, J. E., Wheeler, L., Robbins, I. M., Lane, K. B., Newman, J. H., Johnson, J., Nichols, W. C. & Phillips, J. A., 3RD 2003. Pulmonary veno-occlusive disease caused by an inherited mutation in bone morphogenetic protein receptor II. Am J Respir Crit Care Med, 167, 889-94.

Saito, A., Ochiai, K., Kondo, S., Tsumagari, K., Murakami, T., Cavener, D. R. & Imaizumi, K. 2011. Endoplasmic reticulum stress response mediated by the PERK-eIF2(alpha)-ATF4 pathway is involved in osteoblast differentiation induced by BMP2. J Biol Chem, 286, 4809-18.

Scheuner, D., Song, B., Mcewen, E., Gillespie, P., Saunders, T., Bonner-Weir, S. & Kaufman, R. J. 2001. Translational control is required for the unfolded protein response and in-vivo glucose homeostasis. Mol Cell, 7, 1165-1176.

Sekine, Y., Zyryanova, A., Crespillo-Casado, A., Fischer, P. M., Harding, H. P. & Ron, D. 2015. Stress responses. Mutations in a translation initiation factor identify the target of a memory-enhancing compound. Science, 348, 1027-30.

Shimmi, O., Ralston, A., Blair, S. S. & O’Connor, M. B. 2005. The crossveinless gene encodes a new member of the Twisted gastrulation family of BMP-binding proteins which, with Short gastrulation, promotes BMP signaling in the crossveins of the Drosophila wing. Dev Biol, 282, 70-83.

Shirakawa, K., Maeda, S., Gotoh, T., Hayashi, M., Shinomiya, K., Ehata, S., Nishimura, R., Mori, M., Onozaki, K., Hayashi, H., Uematsu, S., Akira, S., Ogata, E., Miyazono, K. & Imamura, T. 2006. CCAAT/enhancer-binding protein homologous protein (CHOP) regulates osteoblast differentiation. Mol Cell Biol, 26, 6105-16.

Sidrauski, C., Tsai, J. C., Kampmann, M., Hearn, B. R., Vedantham, P., Jaishankar, P., Sokabe, M., Mendez, A. S., Newton, B. W., Tang, E. L., Verschueren, E., Johnson, J. R., Krogan, N. J., Fraser, C. S., Weissman, J. S., Renslo, A. R. & Walter, P. 2015. Pharmacological dimerization and activation of the exchange factor eIF2B antagonizes the integrated stress response. Elife, 4, e07314.

Sonenberg, N. & Hinnebusch, A. G. 2009. Regulation of translation initiation in eukaryotes: mechanisms and biological targets. Cell, 136, 731-45.

Tettweiler, G., Miron, M., Jenkins, M., Sonenberg, N. & Lasko, P. F. 2005. Starvation and oxidative stress resistance in Drosophila are mediated through the eIF4E-binding protein, d4E-BP. Genes Dev, 19, 1840-3.

Thomson, J. R., Machado, R. D., Pauciulo, M. W., Morgan, N. V., Humbert, M., Elliott, G. C., Ward, K., Yacoub, M., Mikhail, G., Rogers, P., Newman, J., Wheeler, L., Higenbottam, T., Gibbs, J. S., Egan, J., Crozier, A., Peacock, A., Allcock, R., Corris, P., Loyd, J. E., Trembath, R. C. & Nichols, W. C. 2000. Sporadic primary pulmonary hypertension is associated with germline mutations of the gene encoding BMPR-II, a receptor member of the TGF-beta family. J Med Genet, 37, 741-5.

Vasudevan, D., Clark, N. K., Sam, J., Cotham, V. C., Ueberheide, B., Marr, M. T., 2ND & Ryoo, H. D. 2017. The GCN2-ATF4 Signaling Pathway Induces 4E-BP to Bias Translation and Boost Antimicrobial Peptide Synthesis in Response to Bacterial Infection. Cell Rep, 21, 2039-2047.

Vattem, K. M. & Wek, R. C. 2004. Reinitiation involving upstream ORFs regulates ATF4 mRNA translation in mammalian cells. Proceedings of the National Academy of Sciences of the United States of America, 101, 11269-11274.

Vazquez de Aldana, C. R., Dever, T. E. & Hinnebusch, A. G. 1993. Mutations in the alpha subunit of eukaryotic translation initiation factor 2 (eIF-2 alpha) that overcome the inhibitory effect of eIF-2 alpha phosphorylation on translation initiation. Proc Natl Acad Sci U S A, 90, 7215-9.

Vuilleumier, R., Springhorn, A., Patterson, L., Koidl, S., Hammerschmidt, M., Affolter, M. & Pyrowolakis, G. 2010. Control of Dpp morphogen signalling by a secreted feedback regulator. Nat Cell Biol, 12, 611-7.

Yang, X., Matsuda, K., Bialek, P., Jacquot, S., Masuoka, H. C., Schinke, T., Li, L., Brancorsini, S., Sassone-Corsi, P., Townes, T. M., Hanauer, A. & Karsenty, G. 2004. ATF4 is a substrate of RSK2 and an essential regulator of osteoblast biology; implication for Coffin-Lowry Syndrome. Cell, 117, 387-98.

Yu, P. B., Beppu, H., Kawai, N., Li, E. & Bloch, K. D. 2005. Bone morphogenetic protein (BMP) type II receptor deletion reveals BMP ligand-specific gain of signaling in pulmonary artery smooth muscle cells. J Biol Chem, 280, 24443-50.

Zid, B. M., Rogers, A. N., Katewa, S. D., Vargas, M. A., Kolipinski, M. C., Lu, T. A., Benzer, S. & Kapahi, P. 2009. 4E-BP extends lifespan upon dietary restriction by enhancing mitochondrial activity in Drosophila. Cell, 139, 149-60.

